# MirGeneDB 2.0: The metazoan microRNA complement

**DOI:** 10.1101/258749

**Authors:** Bastian Fromm, Diana Domanska, Eirik Høye, Vladimir Ovchinnikov, Wenjing Kang, Ernesto Aparicio-Puerta, Morten Johansen, Kjersti Flatmark, Anthony Mathelier, Eivind Hovig, Michael Hackenberg, Marc R. Friedländer, Kevin J. Peterson

## Abstract

Small non-coding RNAs have gained substantial attention due to their roles in animal development and human disorders. Among them, microRNAs are unique because individual gene sequences are conserved across the animal kingdom. In addition, unique and mechanistically well understood features can clearly distinguish *bona fide* miRNAs from the myriad other small RNAs generated by cells. However, making this separation is not a common practice and, thus, not surprisingly, the heterogeneous quality of available miRNA complements has become a major concern in microRNA research. We addressed this by extensively expanding our curated microRNA gene database MirGeneDB to 45 organisms that represent the full taxonomic breadth of Metazoa. By consistently annotating and naming more than 10,900 microRNA genes in these organisms, we show that previous microRNA annotations contained not only many false positives, but surprisingly lacked more than 2,100 *bona fide* microRNAs. Indeed, curated microRNA complements of closely related organisms are very similar and can be used to reconstruct Metazoan evolution. MirGeneDB represents a robust platform for microRNA-based research, providing deeper and more significant insights into the biology and evolution of miRNAs but also biomedical and biomarker research. MirGeneDB is publicly and freely available at http://mirgenedb.org/.

## INTRODUCTION

In the last two decades, the small non-coding RNA field has significantly expanded beyond small nuclear RNAs (snRNAs) and small nucleolar RNAs (snoRNA) (1) to include small interfering RNAs (siRNAs) (2), Piwi-interacting RNA (piRNAs) (3), novel small RNAs derived from known non-coding RNAs, including transfer RNAs (tRNAs), (4) ribosomal RNAs (rRNAs) (5) and microRNAs (miRNAs) (6–9). While each of these types of small RNAs is characterized by a distinctive suite of characteristics and deep evolutionary history, only miRNAs share orthologous gene sequences conserved across Metazoa (10–12). Because of the fundamental roles miRNAs play in establishing robustness of gene regulatory networks across Metazoa (13, 14), and their importance in development (15), formation of cell identity (16) and numerous human diseases including cancer (17, 18), it is imperative that homologous miRNAs in different species are correctly identified, annotated, and named using consistent criteria against the backdrop of numerous other types of coding and non-coding RNA fragments (19–21). Further, it is vital that non-miRNAs are clearly distinguished from *bona fide* miRNAs to avoid spurious conclusions (e.g., (22–24)) concerning the role small RNAs play in human diseases.

Nonetheless, these goals are largely ignored for existing databases, such as miRBase (25), which has developed organically through community-wide submissions of published miRNA calls, and miRCarta (26), a repository that aims to provide miRNA candidates from ultra-deep sequencing experiments in human. With respect to miRBase, several research groups have shown that up to two-thirds of the entries are false positives, with many entries fragments of other classes of small RNAs including tRNAs and snoRNAs, in addition to numerous rRNA fragments (20, 27–36). The interpretation of these non-miRNA fragments as *bona fide* miRNAs affects our understanding of how miRNAs evolve (37), but also incorrectly annotated *bona fide* miRNAs have an impact on the interpretation of data (see (38, 39)). Inconsistencies in nomenclature and changes between releases have made it challenging to use miRBase throughout the years leading to numerous community efforts in extracting information (40–45) and leading to a proliferation of publically available online (see (46)) and study-specific databases (28, 47–51).

To address this, we previously developed a manually curated and open source miRNA gene database, MirGeneDB, which is based on consistent annotation and nomenclature criteria (20). Because it was tightly linked to miRBase and contained only 4 species, the usefulness for comparative studies was heavily limited. Here we present a major update to our database, MirGeneDB version 2.0 (http://mirgenedb.org), which now contains high-quality annotations of more than 10,900 *bona fide* and consistently named miRNAs constituting 1288 miRNA families from 45 species, representing every major bilaterian metazoan group, including many well-established and emerging invertebrate and vertebrate model organisms (Figure 1).

**Figure 1:**
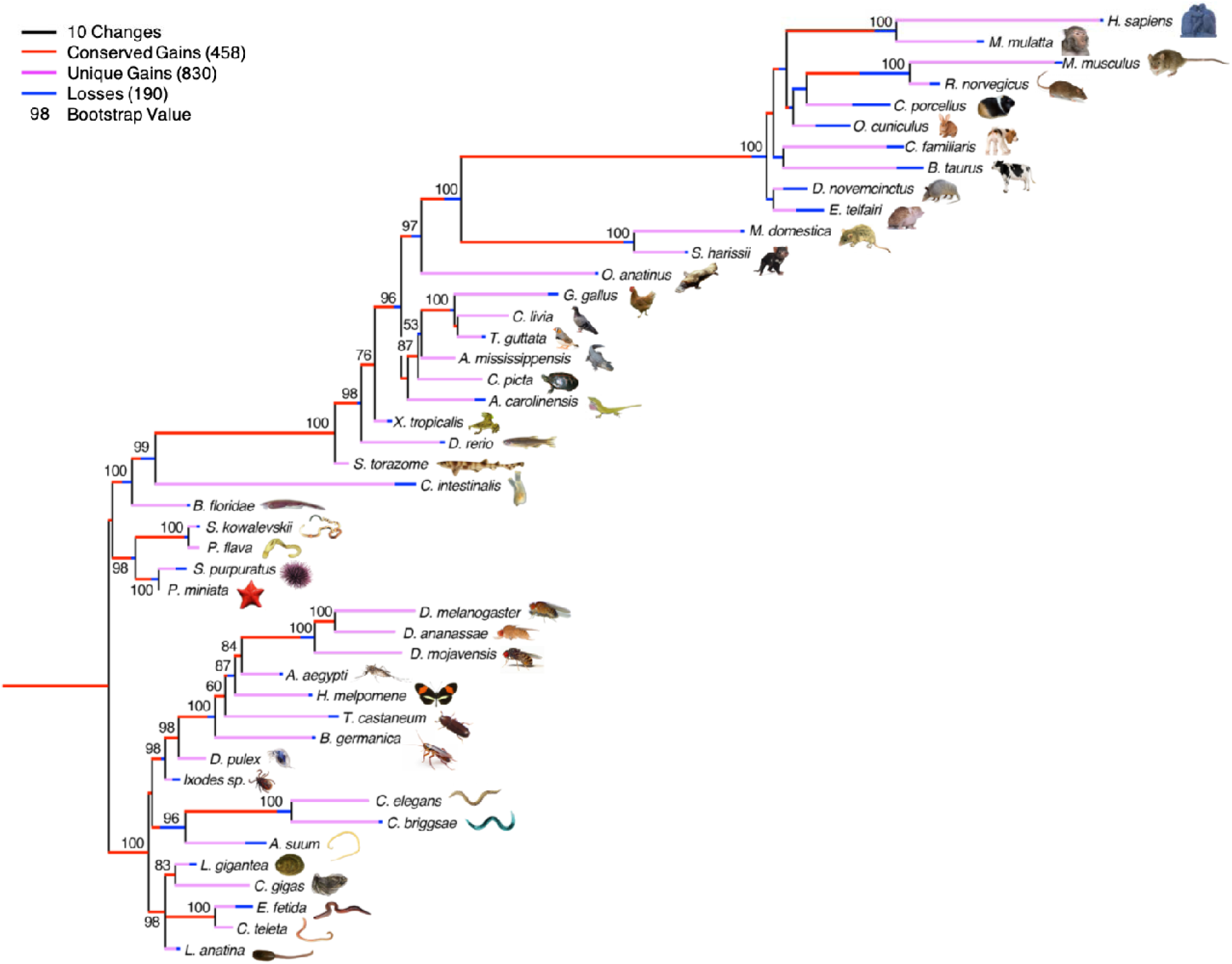
The evolution of the 1288 microRNA families across the 45 bilaterian species currently annotated in MirGeneDB. Conserved gains are shown in red; species-specific gains are shown in pink, and losses are shown in blue; and these gains and losses are mapped onto a generally accepted topology of these species rooted between the deuterostomes and protostomes with branch lengths corresponding to gains and losses, respectively. Note though that this topology is largely recovered when just analyzing the gains and losses of the miRNA families themselves as shown by the bootstrap values indicated at the nodes (Supp. Methods); the only known nodes not recovered are nodes within the placental mammals and Ecdysozoa, primarily due to losses in rodents and nematodes, respectively.

## EXPANSION OF MirGeneDB

For the expansion from version 1.0 to 2.0, we analyzed more than 400 publicly available smallRNA sequencing datasets with at least one representative dataset for each organism, that were automatically downloaded and processed using sRNAbench (52) and miRTrace (53), respectively. This allowed for a consistent and uniform annotation of miRNAomes for each species using MirMiner (11) (see Supplementary Table, “file_info” for files and see Supplementary Information for detailed methods) (20).

The existing MirGeneDB.org miRNA complements for human, mouse, chicken and zebrafish were expanded from our initial effort by 34, 56, 41 and 101 genes to a total of 558, 449, 270 and 388 genes, respectively (Supplementary Table, “table”), and annotation-accuracy for human and zebrafish was further improved using available Cap Analysis of Gene Expression (CAGE) data (Supplementary Figure 1, Supplementary Table, “CAGE”; Supplementary Information) (54). We further used Dicer-, Drosha- and Exportin 5-knockout data (55), as well as primary cell expression data (54, 56–58) to refine human annotations.

Although since its inception MirGeneDB has already given special attention to the annotation of both the 5p and 3p arms, with a clear distinction made between sequenced reads and predicted reads for each miRNA entry, MirGeneDB 2.0 includes four new features related to the transcription and processing of miRNAs (Figure 2A). First, Group 2 miRNAs (59, 60) – those miRNA precursor transcripts that are mono-uridylated at their 3’ end, what we term the 3’ non-templated uridine (3’NTU) – are specifically tabulated, allowing the user to easily discriminate Group 2 from “Group 1” (or canonical) miRNAs. Second, sequence motifs, including the 5’ “UG” motif, the loop “UGU/G” motif, as well as the 3’ CNNC motif (61–63) are bioinformatically identified for every miRNA primary transcript. Third, processing variants, where alternative Drosha/Dicer cuts significantly (>10% of available reads) affect the processed mature seed sequence of the locus (see Manzono et al. 2015, for example (64)), are added as distinct gene entries (indicated with the “v” in the name). Some loci, like the Mir-203 gene (Figure 2A) show both mono-uridylation and variant processing such that only one of the two major variants is classified as a Group 2 miRNA. Curiously, though, in this case the original Drosha cut of Mir-203 results in a proper 2-nucleotide offset for both variants, not a 1-nucleotide offset, and thus both sequences should be properly loaded onto Dicer (60). Therefore, it remains possible that mono-uridylations might serve other or additional purposes beyond Dicer loading as described for some of the vertebrate Let-7s by Kim and colleagues ((60) see below). Finally, we also annotate anti-sense loci (“-as”) for miRNA genes where again significant expression (>10%) of both sense and antisense strands is observed (65).

**Figure 2:**
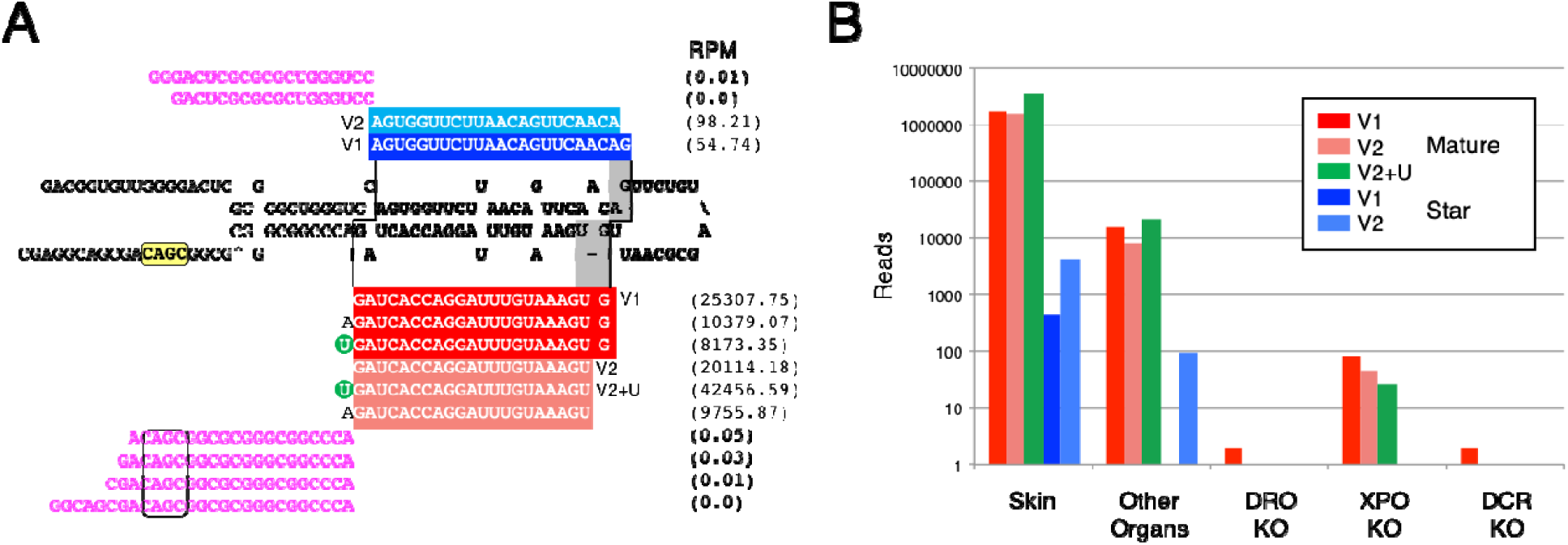
The annotation of microRNA sequences and the implementation of transcriptional and processing information for each miRNA gene in MirGeneDB. **A.** The structure and read stacks for Hsa-Mir-203. The precursor sequence is shown in bold; mature reads are shown in red and star reads in blue with the reads per million for each major transcript detected shown to the far right. A “CNNC” processing motif (63) is shown in yellow. Also shown are the 5’ and 3’ miRNA offset reads (moR) (magenta), which clearly conform to the indicated Drosha cut (staggered line, left) given the reads processed from this locus. The Dicer cut (staggered line, right) results in two primary mature forms (dark vs light red), what we term “variants” (v) that are offset from one another by 1 nucleotide (gray). The 5’ end of variant one starts with the “G” whereas the 5’ end of variant two is moved 1 nucleotide 3’ and starts with the “U.” Each of these two major Dicer products is accompanied by the appropriate star sequence, with variant one shown in dark blue and variant two in light blue. The mature form of variant two – but not version one – is heavily mono-uridylated at its 3’ end (green circle) and is thus a “Group 2” miRNA (55, 60). **B.** The quantification of Hsa-Mir-203 read across various human-specific data sets. As expected (e.g., (66)) expression in skin is about ∼2 orders of magnitude higher relative to other organs sampled (e.g., brain, liver, stomach, lung, uterus, pancreas, testes, colorectum, small intestine and kidney) and the detection of the mature form is nearly 3 orders of magnitude relative to the star. Consistent with Mir-203 being a *bona fide* miRNA, expression is nearly abrogated in DROSHA and DICER knock-out, and greatly diminished in the EXPORTIN-5 knock-out (55).

### QUALITY OF MirGeneDB ANNOTATIONS

Any database should be largely free of both false positives and false negatives. To account for this, miRBase categorizes a subset of their entries as high-confidence miRNAs, which are those that are highly expressed and show clear indications of proper processing, and further has introduced a public voting system to identify more high-quality candidates (67). MirGeneDB takes an alternative approach: rather than allowing for community annotation, the near-complete miRNA repertoire of each taxon is added to MirGeneDB using a consistent and well-defined set of criteria (19, 20, 68). When comparing MirGeneDB 2.0 and miRBase, the number of miRNAs conforming to the annotation criteria is about 3 times higher in MirGeneDB than it is in miRBase (2,844 for the miRBase ‘high confidence’ set (67)). Further, because the primary requirement for the inclusion of a putative miRNA to miRBase is publication in a peer-reviewed journal, over time, miRBase has become increasingly heterogeneous with respect to the number of miRNAs for closely related species, such as the often studied human and rarely studied macaque (69) (Fig 3.). This focus on model systems – in particular human, mouse and chicken – has resulted in miRBase having, on the one hand, a much larger number of annotated sequences for some of the 38 taxa shared with MirGeneDB2.0, accounting for estimated 5,617 false positives, and, on the other hand, miRBase lacking 22% of all MirGeneDB2.0 genes, accounting for 2,118 false negatives (Supplementary Figure 2, Supplementary Table, “overview”). These disparities have obstructed comparative genomic approaches in the miRNA field: for example, missing miRNA families have been misinterpreted as secondary losses, questioning then the fundamental conservation of miRNA families (70). However, very similar miRNA complements in terms of total miRNA genes and miRNA families are observed in closely related groups in MirGeneDB (Figure 3), supporting earlier evolutionary studies arguing for the utility of miRNAs as excellent phylogenetic markers (11, 37, 53, 68) (Figure 1).

**Figure 3:**
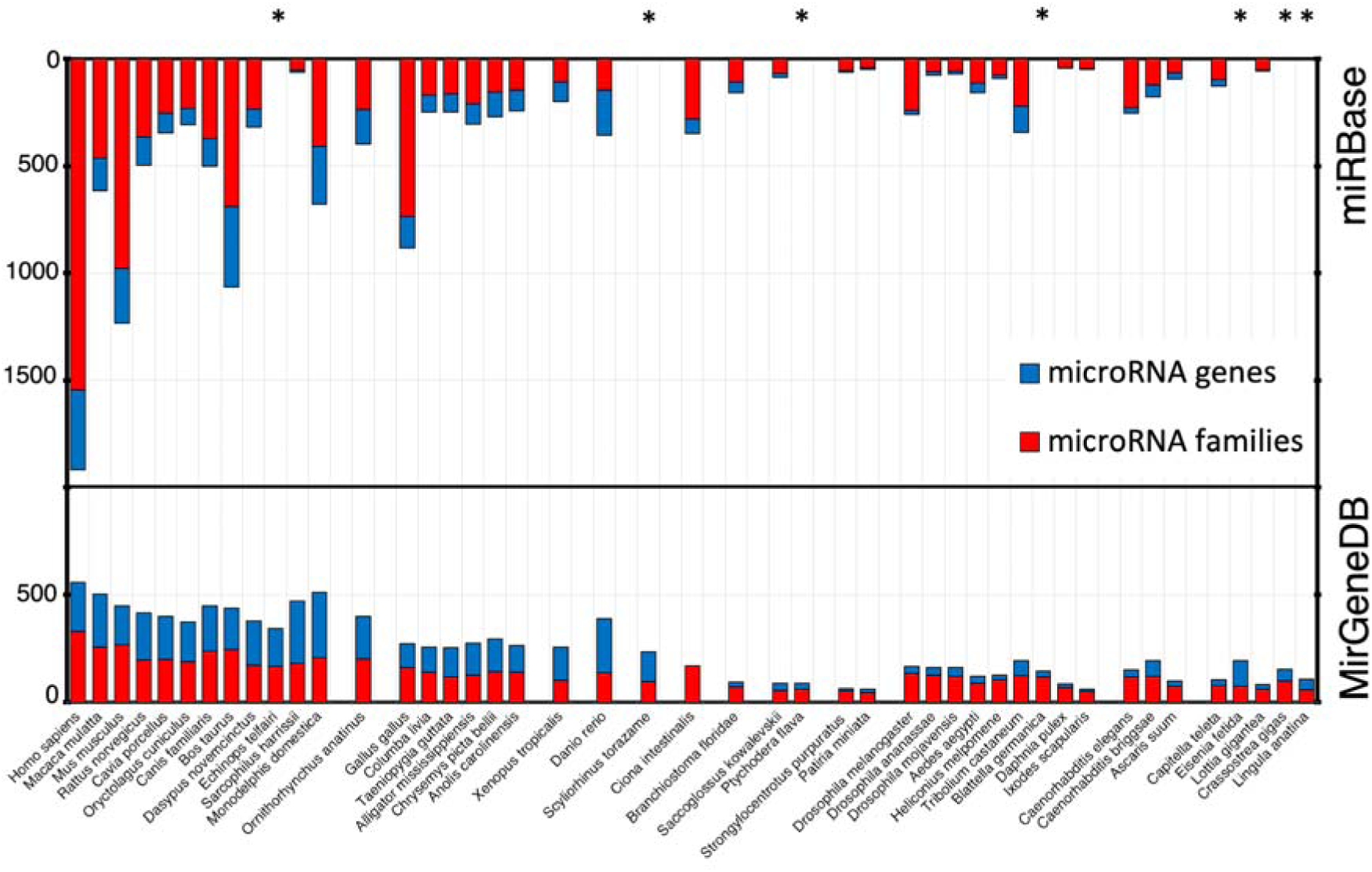
Metazoan miRNA complements are homogeneous between closely related species. Top: miRBase community-report based complements show high heterogeneity in the numbers of families (red) and genes (blue) for closely related species. For instance, in miRBase, human and macaque differ by 1300 genes (Hsa 1917, Mml 617) and 1081 families (Hsa: 1543, Mml: 462). Bottom: MirGeneDBs curated complements are homogeneous for both gene and family numbers (see Supplementary Figure 3 for conserved families, genes in comparison to novel families and genes). For instance, in MirGeneDB, human and macaque differ by 56 genes (Hsa 558, Mml 502) and 38 conserved families (Hsa: 267, Mml: 229). Asterisks mark species that are found in MirGeneDB, but not in miRBase.

Thus, while it is inevitable that some cell-type specific or lowly expressed miRNAs are missing from our annotations, MirGeneDB can be considered essentially free of false positives. Further, because MirGeneDB is focused on identification of miRNA genes and families, rather than sequences (20), a *bona fide* miRNA gene identified in one taxon is identified as such in all, in contrast to miRBase, where the same gene can be identified as generating a high-confidence miRNA sequence in one taxon, but a low-confidence sequence in another (20). Hence, we are confident that there are few (if any) missing miRNA genes that are conserved between two (or more) of the 45 currently included taxa.

## IMPROVED WEB INTERFACE OF MIRGENEDB

The expanded web-interface of MirGeneDB 2.0 allows browsing (http://mirgenedb.org/browse), searching (http://mirgenedb.org/search) and now also downloading (http://mirgenedb.org/download) of miRNA-complements for each organism, in addition to a general information page about the criteria used for miRNA annotation (http://mirgenedb.org/information), as well as false negatives for each taxon (where known), with links to previous versions of MirGeneDB. On the *browse-pages* for each organism (e.g. http://mirgenedb.org/browse/hsa), a table is available that includes MirGeneDB names and miRBase names (if available), family-level assignment and the strandedness of the miRNA (i.e., whether the mature arm is the 5p arm, the 3p arm, or both) (Figure 4, “A”); overview information on location in the genome (Figure 4, “B”); and the phylogenetic origin of each miRNA locus and family (Figure 4, “C”). The new features in MirGeneDB 2.0, including the 3’ NTU’s and sequence motifs (see Figure 2) are also indicated (Figure 4, “D”). Finally, a heatmap of the expression of each miRNA for all available tissues is available to orient users on expression patterns (Figure 4, “E”).

**Figure 4:**
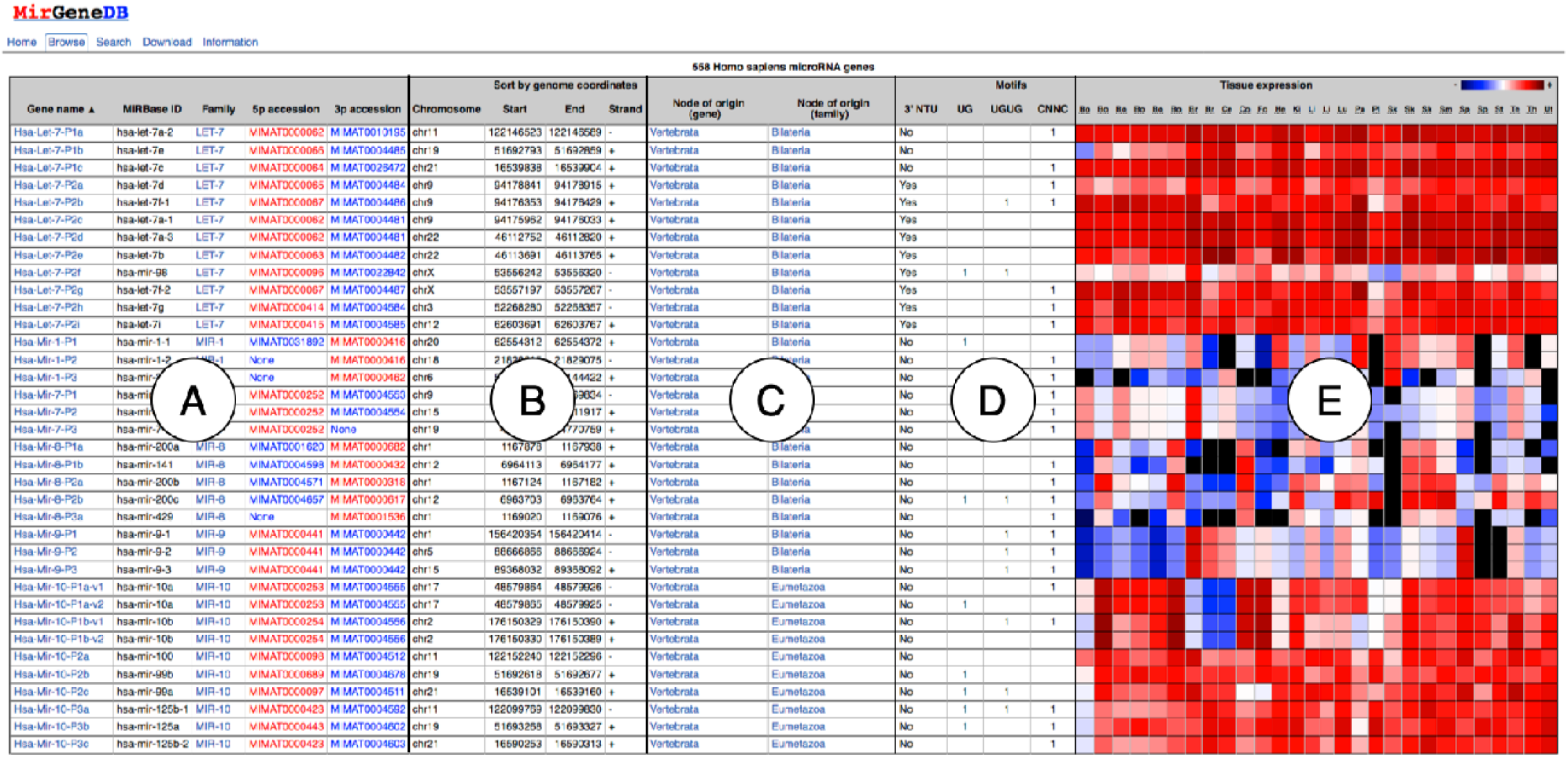
Improved web interface of MirGeneDB. For each species in MirGeneDB an overview browse page exists that lists all genes. For each gene the following information is provided and sortable: names (both MirGeneDB and miRBase), family-level assignments, and strandedness (A), genomic coordinates (B); inferred phylogenetic origin of both the gene locus and family (C); information on the presence or absence of 3’ NTU’s and sequence motifs (D); and a normalized heatmap for available datasets.

From here, *gene-pages* for each miRNA gene can be opened that contain names, orthologues and paralogues, sequences, such as the mature seeds, structure, and a range of other information, including genomic coordinates (e.g. http://mirgenedb.org/show/hsa/Let-7-P1a). Further, interactive *read-pages* are also provided for each gene (e.g., http://mirgenedb.org/static/graph/hsa/results/Hsa-Let-7-P1a.html) that show an overview of read-stacks on the corresponding extended precursor sequence of each *gene-page.* These pages contain detailed representations of templated and 3’-end non-templated reads for individual datasets for each gene, including reports on miRNA isoforms and downloadable read-mappings, and the information can be used to quantify expression of any miRNA across known data sets (e.g., Figure 2B).

We provide highly comparative data with distinct and accurate sub-annotations of the precursor, mature, loop, co-mature or star sequences. In addition, we also provide 30-nucleotide flanking regions on both arms for each miRNA to generate an extended precursor transcript for the discovery of regulatory sequence motifs, and lastly seed sequences are also annotated separately. On the *search-pages*, all these annotations can be searched independently, either by sequence using Blast (71), or, if existing, by the full miRBase name (72), and by the MirGeneDB name, following our consistent nomenclature (20). Users can also search specific 7-nt seed sequences, and all searches can be done either for individual species or over the entire database. Finally, on the *download-pages*, fasta, gff, or bed-files for all miRNA components are downloadable for each species.

## MiRNA NOMENCLATURE

Following Ambros et al. (19), MirGeneDB 2.0 employs an internally consistent nomenclature system where genes of common descent are assigned the same miRNA family name, allowing for the easy recognition of both orthologues in other species, and paralogues within the same species, as described earlier (20).

It was recently argued that miRNA classification should reflect the similarity of the seed, and hence similarity of function, which corresponds their targeting properties (63). However, as Darwin recognized, *natural* classification systems are never functional, instead “… all true classification is genealogical.” (73). Classifying biological entities by function rather than evolutionary history risks creating confusion as Ambros et al. (19) recognized, and thus a proper understanding of both miRNA form and function requires recognizing and describing miRNAs within the context of their unique (but not necessarily entirely discoverable) evolutionary history. However, we have taken steps to accommodate the functionally focussed community (see below).

The advantages – and limitations – of the nomenclature system employed by MirGeneDB are exemplified by the LET-7 family of miRNAs (Figure 5). Let-7 is an ancient miRNA gene evolving sometime after the bilaterian split from cnidarians, but before the divergence between protostomes and deuterostomes, and was (and, in many taxa, still is) syntenically linked to two MIR-10 family members. However, before the last common ancestor of urochordates and vertebrates (collectively called the Olfactores, (74)), this original gene duplicated, generating two paralogues, one still being linked to the two MIR-10 genes (paralogue 1, light gray box), and a second, now located elsewhere in the genome (paralogue 2, dark gray box) (75), that is mono-uridylated at the 3’ end (60). This second paralogue duplicated several times before the divergence between urochordates and vertebrates, and then, early in vertebrate evolution, the entire genome duplicated twice, generating ancestrally 14 Let-7 paralogues in the last common ancestor of gnathostomes, three derived from the initial Let-7 gene (named P1a-c) and 11 (named P2a-k) from the second. However, the duplication history of these 11 vertebrate P2 paralogues cannot currently be completely elucidated (see also (75)), although the orthology can, and hence they are simply named P2a-k. Further, these paralogues cannot be orthologized with the five P2 genes in urochordates, so these genes are called “orphans” (20) in the urochordate, to highlight the fact that they cannot be currently conclusively orthologized with vertebrate P2 genes. However, if new information comes to light that will allow for robust phylogenetic insight, these names would be changed accordingly.

**Figure 5.**
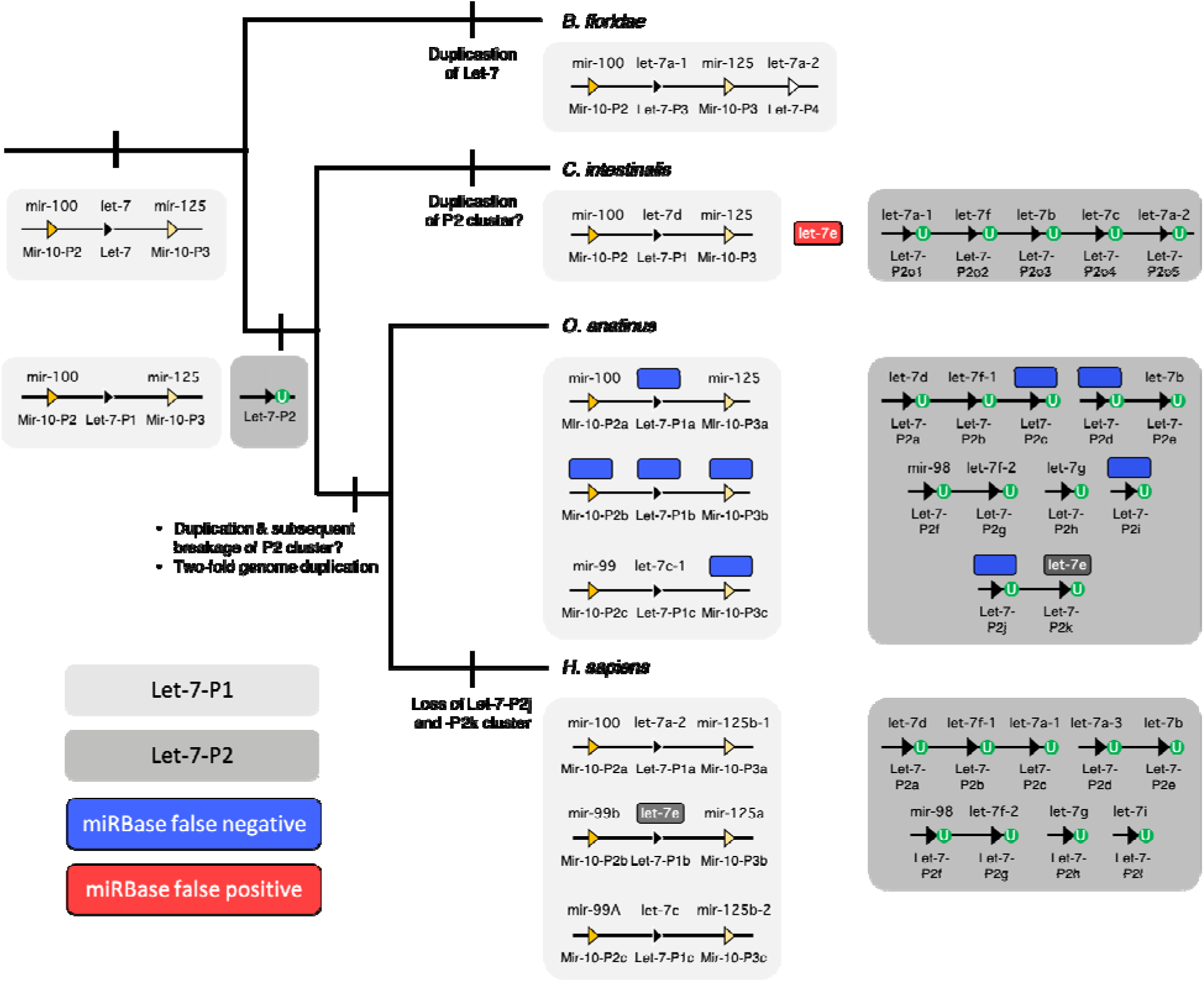
Nomenclature comparison between MirGeneDB and miRBase for representative chordate Let-7s. Shown is the accepted topology (74) for the three major subgroups of chordates, and for each taxon, a (unscaled) representation of the genomic organization of its Let-7 genes/sequences. MirGeneDB names are shown below each of the loci symbols, and the miRBase sequence names are above. The primitive condition is to possess a single Let-7 gene linked to the two Mir-10 genes (light gray box), as is still found in many bilaterian taxa. In the amphioxus *Branchiostoma floridae*, this single Let-7 duplicated, and this new paralogue is now positioned at the 3’ end of the cluster. In the Olfactores there is a separate gene duplication event generating another paralogue that is not linked to the original Let-7 cluster in any known urochordate, like *Ciona intestinalis* or any vertebrate, including human (*H. sapiens*) or the platypus (*O. anatinus*). Further distinguishing this paralogue is the fact that in all Olfactores these Let-7 genes (shown in the dark gray boxes) are Group 2 miRNAs, each with an untemplated mono-uridylated 3’ end (green circles) (see (60)). False negatives (i.e., loci present and transcribed that are present in MirGeneDB, but not in miRBase) are shown in blue. A single false positive (i.e., a sequence present in miRBase – cin-let-7e – but without a corresponding locus in the genome) is shown in red. Note that let-7e also names two sequences derived from two non-orthologous genes in human and platypus – a canonical Group 1 Let-7 (Let-7-P1b) in human, but a Group 2 miRNA (Let-7-P2k) in platypus. This locus is also in diapsids (birds and ‘reptiles’), as well as in the teleost fish *Danio rerio*, but is lost in therian (i.e., placental and marsupial) mammals (see also Hertel et al. 2012). Despite the fact that the monophyly of these Group 2 Let-7s in Olfactores appears robust, how these ancestral 11 genes in bony fishes are related to one another, and how they relate to the five genes in *C. intestinalis*, remains unknown given the lack in phylogenetic signal in the primary precursor sequences themselves and the apparent absence of syntenic signal as well. Hence, MirGeneDB names these genes with this phylogenetic opacity in mind.

The nomenclature advantages of MirGeneDB over miRBase are obvious. First, non-orthologous genes are never given the same name. For example, both human and platypus have let-7e sequences, but let-7e in human is derived from the ancestral P1 gene is linked to MIR-10 genes, and is a Group 1 miRNA; let-7e in platypus is derived from the ancestral P2 gene, is not linked to MIR-10 genes, is mono-uridylated at its 3’ terminus, and maybe most importantly is a gene lost in all therian (i.e., placental and marsupial) mammals (Figure 5).

Second, simply from the name, one can get an accurate picture of the evolutionary history of the gene within the context of a monophyletic miRNA family (20). For example, there are two Let-7 genes in the amphioxus *Branchiostoma floridae*, a close chordate relative to urochordates and vertebrates. How are these genes related to the ancestral gene duplicates in Olfactores? miRBase names them let-7a-1 and let-7a-2, the same names employed by two human miRNAs. However, these two genes are amphioxus-specific gene duplicates of the MIR-10 associated Let-7 gene, and are thus named Let-7-P3 and Let-7-P4 to distinguish these unique paralogues from the two Let-7 paralogues (P1 and P2) of Olfactores (Figure 5).

Third, misnamed genes will not be orphaned in literature searches or functional studies. For example, one of the 12 human Let-7 paralogue was originally named mir-98 (see Figure 5), and although miRBase lists this gene correctly within the LET-7 family, it is not obvious from the name itself. Notably, in the latest release, applying a novel text mining approach for literature searches, the miRBase authors state that there are only 11 Let-7 family members in human, failing to account for Mir-98 (25). This example clearly highlights the importance of consistent naming and the risks of non-uniform nomenclature systems.

Finally, because MirGeneDB uses this natural classification and nomenclature system, it allows for an accurate reconstruction of ancestral miRNA repertoires – both at the family-level and at the gene--level – that is now provided in MirGeneDB 2.0 for all nodes leading to the 45 terminal taxa considered. This allows users to easily assess both gains and losses of miRNA genes and families through time. Again, with respect to the LET-7 family, it is clear that therians lost two ancestral Let-7 genes, genes that are still retained in platypus (Let-7-P2j and -P2k, see Figure 5) and were present in the last common ancestor of all living bony fishes. In order to not increase confusion about the naming of miRNA genes though, we continue to provide commonly used miRBase names – if available – in our “*browse*” section of MirGeneDB2.0 for miRNA sequences (e.g., http://mirgenedb.org/browse/hsa, see Figure 4 column “A”).

Nonetheless, in order to fit the needs of the scientific community focussed on miRNA seeds and their targets, we have introduced “seed-sequence” entries for each mature miRNA that summarize all entries with the exact same seed sequence (see for instance http://mirgenedb.org/browse/ALL?seed=GAGGUAG, and note that 4 paralogues of the 3’ mature sequences of MIR-7594 in *C. briggsae* share the seed with 305 5’ mature Let-7 sequences in all 45 MirGeneDB species), which will allow, together with our seed-blast function, for easy lifting to the functional seed-based definition for individual miRNAs.

## FUTURE DEVELOPMENTS

The establishment of this carefully curated database of miRNA genes, supplementing existing databases, including miRBase and miRCarta, represents a stable and robust foundation for reproducible miRNA research, in particular studies that rely on cross-species comparisons to explore the roles miRNAs play in development and disease, as well as the evolution of miRNAs and animals themselves. Our long-term goal is to have a wider representation of bilaterian metazoan species, and for each of these organisms a large number of comparable datasets for a comprehensive set of organs, tissues and cell types.

We hasten to stress that although all nearly 11,000 genes currently in MirGeneDB have been hand curated, mistakes are inevitable, both in terms of the inclusion of species-specific false positives, missing false negatives, as well as processing errors, mistakes in understanding evolutionary history (possibly resulting in nomenclature errors), and other factors. We would ask the community to alert us to any such errors as only through community-wide collaboration can these inevitable mistakes be eliminated from the database, and MirGeneDB promises to resolve any errors in a timely fashion.

## DATA AVAILABILITY

All MirGeneDB data are publicly and freely available under the Creative Commons Zero license. Data are available for bulk download from http://mirgenedb.org/download. Feedback on any aspect of the miRBase database is welcome by email to, or via Twitter (@MirGeneDB).

## ACKNOWLEDGMENTS

We thank Marc Halushka and Gianvito Urgese for discussions, and Georgios Magklaras and Sveinung Gundersen for IT support.

## FUNDING

BF, WK and MRF acknowledge funding from the Strategic Research Area (SFO) program of the Swedish Research Council (VR) through Stockholm University. BF and EH were supported by South-Eastern Norway Regional Health Authority grant #2014041 and #2018014, respectively. VO would like to acknowledge Russian Science Foundation (grant number 18-15-00098), Dr Mary J. O’Connell and the School of Life Sciences at the University of Nottingham for supporting this work. AM has been supported by the Norwegian Research Council, South-Eastern Norway Regional Health Authority, and the University of Oslo through the Centre for Molecular Medicine Norway (NCMM), which is part of the Nordic European Molecular Biology Laboratory partnership for Molecular Medicine. K.J.P. has been supported by the National Science Foundation, NASA-Ames, and Dartmouth College.

## Notes

#### Summary of Updates

This is the final version of MirGeneDB 2.0 and it has been submitted for publication. 45 species and nearly 11,000 genes in total.

http://mirgenedb.org/

