## Supplementary Information for "MirGeneDB 2.0: The metazoan microRNA complement"

***Features of miRNAs***

MiRNAs can be distinguished from other small RNA families by a set of unique features as described earlier (1, 2): The presence of two 20-26nt long reads that are expressed from each of the two arms derived from a stable hairpin precursor is essential to assess whether or not Drosha and Dicer were involved in the processing. Since the ends of canonical miRNA reads are generated enzymatically, the 5’ ends of the reads are homogeneous (>90%). The hairpin precursor shows imperfect complementarity and base pairs in at least 16 of the ~ 22 nucleotides. The 5p and 3p reads are offset by 2 nucleotides on both ends due to the sequential processing of the miRNA transcript by Drosha and Dicer to generate the mature ~22 nucleotide read(s). In some cases, the Drosha offset is only offset by 1 templated nucleotide, but in these cases the 3’ end of the 3p arm is monouridilyated (3, 4). The length of the loop is at least 8 nucleotides long; there is no apparent maximum in loop length, even in organisms possessing only a single Dicer gene, contra our earlier statement (2), even though most taxa like vertebrates with single Dicer genes never show loop lengths greater than ~40 nucleotides.

There are other features of miRNAs, in particular structural and evolutionary signatures that allow them to be further distinguished from other small RNAs. The mature miRNA sequence usually starts with A or U, and is often mismatched with the complementary arm, which seems to facilitate arm selection by Argonaute (at least in mammals) (2, 5, 6). Nucleotide positions 2 through 8 of the mature sequence (the "seed") are strongly conserved through evolution, as are positions 13-16 (the 3' complementary region) (2, 7). Recently it was demonstrated that processing motifs are often (but not always) present in the primary miRNA transcript including a UG motif 14 nucleotides upstream of the 5p arm, a UGU motif at the 3' end of the 5p arm, and a CNNC motif 17 nucleotides downstream of the 3p arm (8–10).

***Processing of sequencing data and expression profiling***

Publicly available smallRNA sequencing data of whole organisms, healthy organs, tissue or cell-isolates was downloaded from European Genome-phenome Archive (EGA), the Sequence Read Archive (SRA) and the Gene Expression Omnibus (GEO) respectively (see Supplementary Table, “file_info”). For download and processing we used the latest version of sRNAbench (11) and miRTrace (12), respectively. Corresponding files were automatically downloaded and converted into fastq files. All datasets were consistently processed. Briefly, i) : 3' adapter sequences were automatically identified using sRNAbench and processed by miRTrace (“qc mode”) (11), ii) reads containing ambiguous bases (N) were eliminated, iii) reads < 18nt and > 27 nt were filtered out, iv) remaining reads were collapsed into unique read entries (sequence & read count). Collapsed reads were mapped to MirGeneDB (2) using bowtie1.2 (13), requiring an 18 nucleotide seed sequence of zero mismatches to avoid cross-mapping. All mappings were transformed to bam-files using SAMtools (14).

***Refinement of pre-miRNA 3’end annotation with CAGE data***

Human annotation

We downloaded the hg38 bigwig files associated to all ENCODE CAGE experiments from the ENCODE data portal (see https://www.encodeproject.org/metadata/type=Experiment&assay_slims=Transcription&assay_title=CAGE&assembly=GRCh38&files.file_type=bigWig/metadata.tsv). We merged the data from all the experiments and converted the files in the BED format. Computation and plotting of the distribution of CAGE tags around the 3’ end of pre-miRNAs annotated in MirGeneDB were performed using the deepTools v. 2.4.0 (Supplementary Figure 1a). As described in previous studies (15, 16) we observed a peak for CAGE tags 1 nt downstream (i.e. the +1 nt) of the pre-miRNA 3’ ends (Supplementary Figure 1). We considered for manual curation the pre-miRNAs showing a higher number of CAGE tags at positions 0 or +2 with respect to the annotated pre-miRNA 3’ end, which could correspond to a 1 nt off misannotation (see<http://fantom.gsc.riken.jp/zenbu/gLyphs/#config=BIP45GpJGl7o5GEvt0umIC> for an example). After manual curation through the Zenbu genome browser (17), we corrected the 3’ end position for pre-miRNAs Hsa-Mir-145, Hsa-Let-7-P2i, Hsa-Let-7-P2d (Supplementary Table, “CAGE”).

Zebrafish annotation

We applied the same methodology to the CAGE data obtained from 12 developmental stages of embryogenesis in zebrafish (15). Bigwig files of CAGE tags mapping were retrieved using the CAGEr R package (18). Data from all developmental stages were merged to analyze the distribution of CAGE tags around pre-miRNA 3’ ends (Supplementary Figure 1b). After manual curation, we updated the 3’ end position of the pre-miRNAs Dre-Mir-153-P1a and Dre-Let-7-P2c (Supplementary Table, “CAGE”).

Water flea annotation

CAGE data for *Daphnia pule*x derived from three developmental states were retrieved from GEO (GSE80141) (19). We followed the same steps as described above for human and zebrafish CAGE data but did not find any 3’ end annotation of pre-miRNAs to update.

***Phylogenetic analysis***

A data matrix of 1288 miRNA families scored for 45 taxa (available upon request) was analyzed with PAUP v. 4.0a (build 165) for Macintosh (20). Parsimony analysis used Dollo parsimony and the bootstrap analysis used a full heuristic search with a 1000 replicates.
