## Supplementary Figures for "MirGeneDB 2.0: The metazoan microRNA complement"

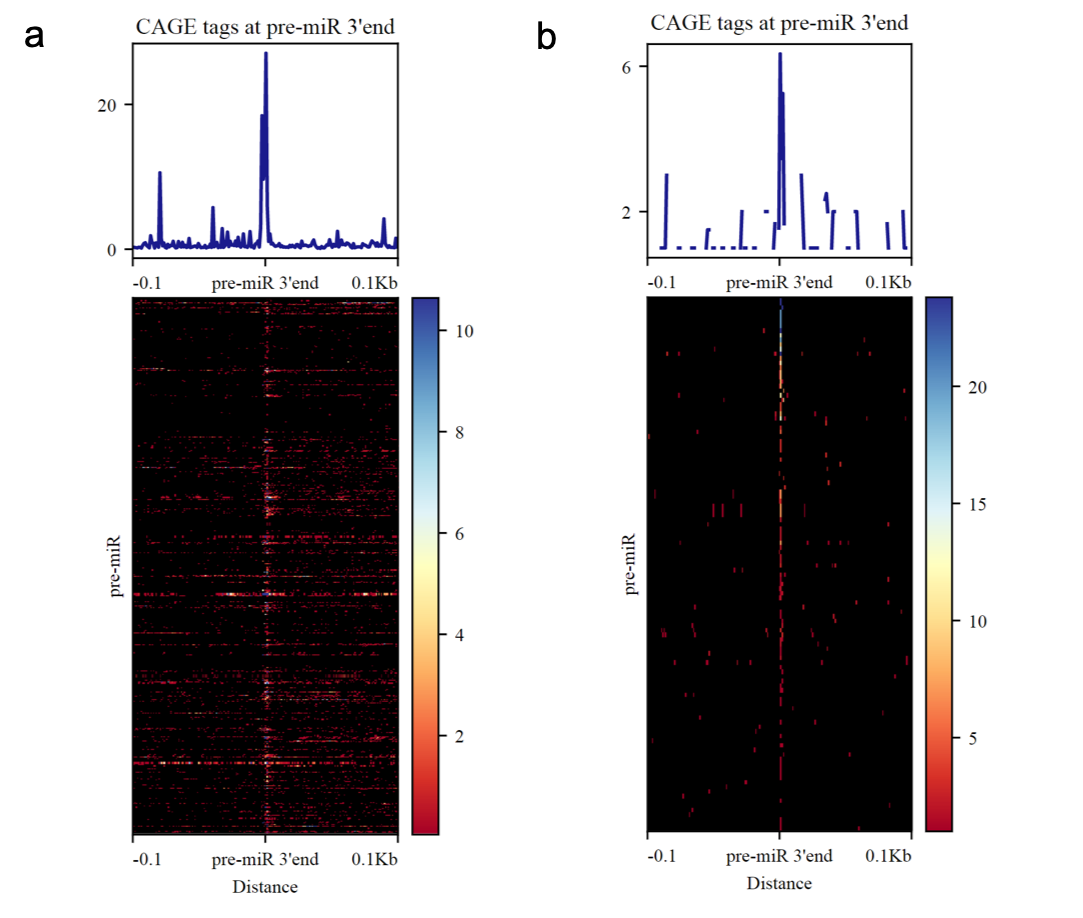

**Supplementary Figure 1**: The distribution of CAGE tags around the 3’ end of pre-miRNAs annotated 329 in MirGeneDB for a) human and b) zebrafish shows a clear peak for CAGE tags 1 nt downstream (i.e. the +1 nt) of the pre-miRNA 3’ ends as described before (1, 2).

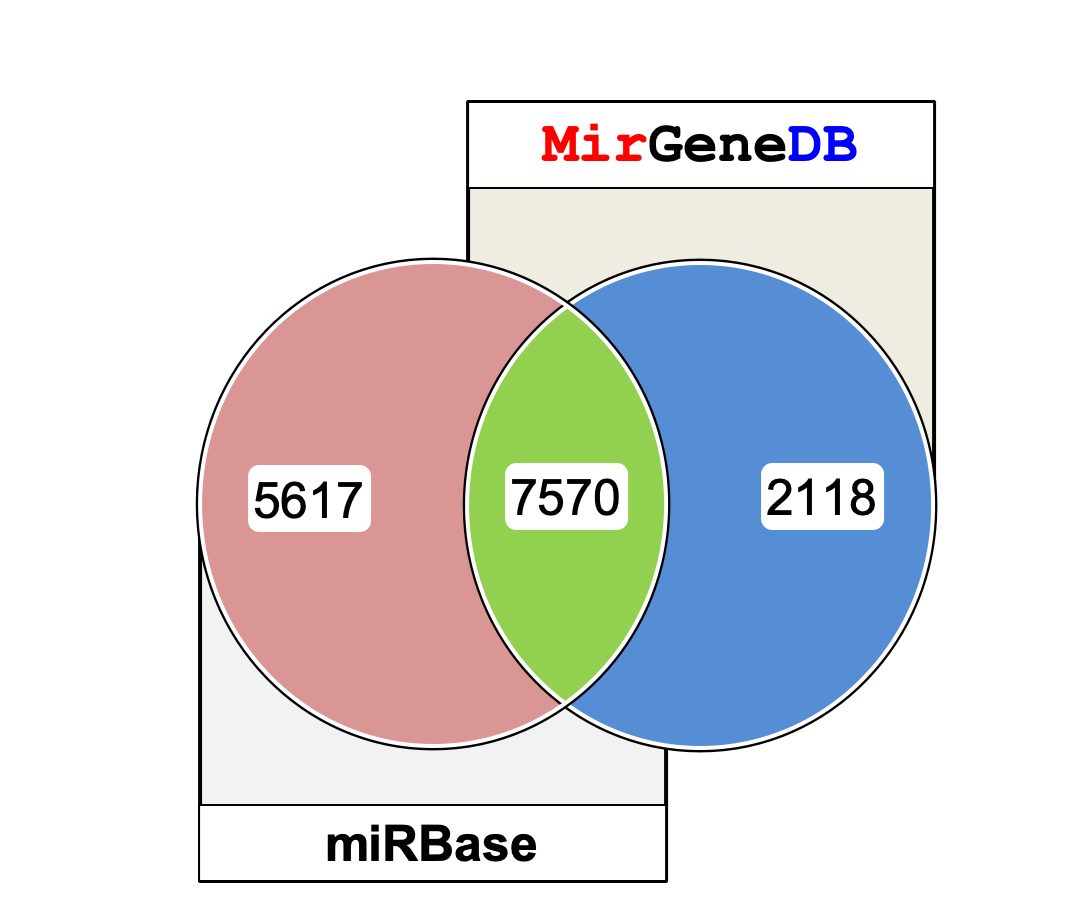

**Supplementary Figure 2:** The comparison of the microRNA complements of 38 organisms shared between miRBase and MirGeneDB revealed that only 7,570 of the 13,187 entries in miRBase where common with MirGeneDB (green) and 5,617 entries represented *false positive* entries (red). Additional 2,118 miRNA genes that were annotated in MirGeneDB 2.0 for the 38 species were not found in miRBase (= *false negatives*) (blue).

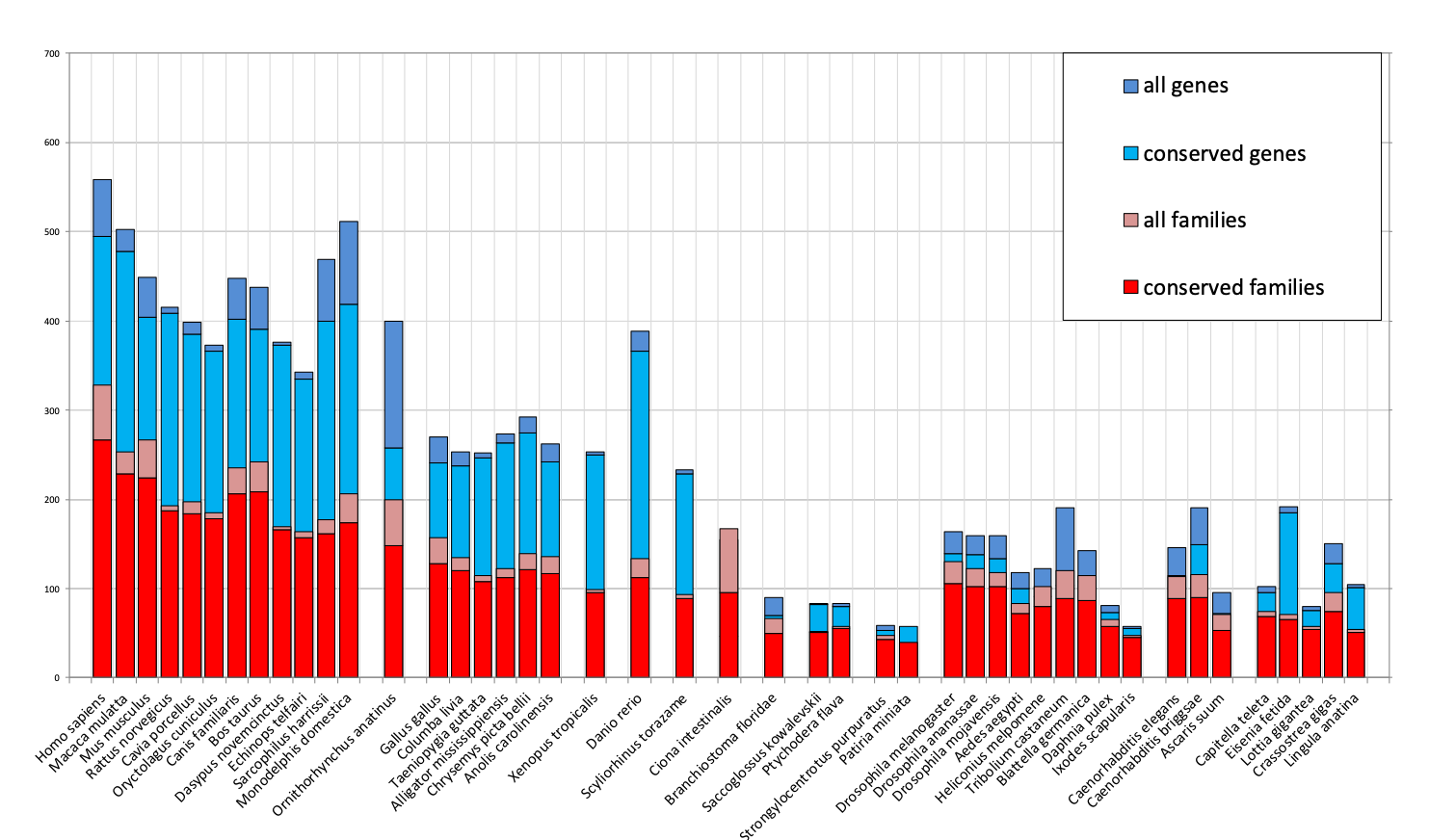

**Supplementary Figure 3**: High congruence of conserved miRNA families and genes between closely related organisms. Conserved miRNA families (red) show the highest congruence between closely related organisms, followed by conserved genes (light blue).
